# Demonstration of SLU7 as a new pan-cancer target

**DOI:** 10.1101/2025.08.25.672085

**Authors:** C. Rojo, A. Otero, M. Elizalde, M. Azkona, R. Barbero, MU. Latasa, I. Uriarte, A. Gutierrez-Uzquiza, D. Alignani, L Guembe, A. Lujambio, F. Pastor, MG. Fernández-Barrena, MA. Ávila, M. Arechederra, C. Berasain

## Abstract

Cancer treatment remains challenging due to heterogeneous responses to immunotherapy across patients and tumor types. Innovative strategies are required to overcome immune evasion. We have identified the splicing factor SLU7 as essential for the survival of cancer cells from diverse origins. SLU7 knockdown induces R-loop accumulation, transcription-dependent genomic instability, DNA damage, and replication catastrophe, together with aberrant splicing and inhibition of nonsense-mediated mRNA decay (NMD) and/or DNA methylation. These alterations lead to the expression of neoantigens, interferon B1, endogenous retroviruses, and cancer-testis antigens, which would enhance tumor immunogenicity.

Therefore, we propose SLU7 targeting as a dual-action therapy, combining direct tumor suppression with immune activation. Using various murine cancer models, including orthotopic liver tumors, and multiple molecular strategies—such as inducible CRISPR/Cas9, systemic delivery of chimeric siSLU7–nucleolin aptamers (APTASLU), and intratumoral injection of siSLU7-loaded nanoparticles—we show that distinct siSLU7 sequences and delivery platforms effectively inhibit tumor growth. Furthermore, SLU7 silencing synergizes with immune checkpoint inhibitors, amplifying anti-tumor responses.

Our *in vivo* data demonstrate that SLU7 is a promising, versatile target for diverse cancers. Its multimodal mechanism offers potential to overcome tumor heterogeneity, reverse immune tolerance, and enhance immunotherapy efficacy.

## Introduction

Immunotherapy —particularly through the use of immune checkpoint inhibitors (ICIs)— has transformed the treatment landscape for patients with solid tumors, including hepatocellular carcinoma (HCC), one of the most prevalent cancers globally and the third leading cause of cancer-related death worldwide^1^. Despite this progress, a substantial proportion of patients fail to respond to ICIs, largely due to diverse immune evasion mechanisms, such as the inherently low immunogenicity of certain tumors^1,2^. This underscores the urgent need for novel strategies to boost anti-tumor immune responses^2^.

To address this challenge, several approaches are under investigation. Among them, epigenetic therapies such as DNMT1 inhibitors^3^, splicing modulators^4^, and nonsense-mediated decay (NMD) inhibitors^5^ have shown promise in enhancing the efficacy of ICIs in both preclinical models and early clinical trials.

Our previous studies identified the splicing factor SLU7 as a critical survival factor in various cancer cell types, including HCC, cervical carcinoma, and non-small cell lung cancer (NSCLC)^6^. We further demonstrated that SLU7 knockdown induces apoptosis across these cancer types, preceded by a cascade of molecular events: oxidative stress^7^, R-loop accumulation^8^, transcription-dependent genomic instability^8^, DNA damage^8^, replicative catastrophe^8^, inhibition of DNA methylation^9^, and disruption of NMD^10^. These alterations lead to the generation of aberrant splicing and NMD isoforms, derepression of antigens normally silenced by DNA methylation, transcriptional dysregulation, and reduced protein stability^11^.

Based on these findings, we hypothesized that SLU7 downregulation could serve as a novel anti-cancer strategy both as monotherapy and in combination with ICIs, by enhancing tumor antigenicity and immune recognition.

In this study, we demonstrate that the dependency on SLU7 for cell survival is a widespread vulnerability across multiple cancer types, including HCC, NSCLC, melanoma, colon, breast, cervical carcinoma, and cholangiocarcinoma (CCA). We conducted proof-of-concept experiments showing that intratumoral gene editing or downregulation of SLU7 expression significantly impairs tumor growth. Furthermore, we evaluated various therapeutic strategies—including aptamer-siSLU7 chimeras and magnetic nanoparticles conjugated with siSLU7 oligonucleotides—in both heterotopic and orthotopic syngeneic and xenograft models. Our findings indicate that SLU7 silencing represents a promising approach to cancer therapy, with potential to significantly enhance the efficacy of immunotherapy.

## Results

### 1. SLU7 is a survival factor for tumor cells

We have previously demonstrated that SLU7 is a survival factor for human HCC cells^6^. We found that while SLU7 silencing does not affect the survival of normal hepatocytes it induces the apoptotic death not only of HCC cells but also of HeLa cervical carcinoma cells and H358 NSCLC cells^6^.

Our present results demonstrate that a large variety of transformed cells depend on SLU7 for survival. As shown in **Figure 1A**, SLU7 silencing induces DNA damage, as evidenced by γH2AX increase, and caspases activation, as evidenced by PARP1 cleavage, resulting in cell death (**Figure S1A**) of human melanoma, colon cancer, NSCLC and cholangiocarcinoma (CCA) cell lines. Furthermore, mouse melanoma, breast cancer and HCC cell lines are also dependent on SLU7 for survival (**Figure S1** and **Figure 1B**). Moreover, consistent with the results observed in human cell lines, and the impact of SLU7 downregulation on splicing, NMD inhibition, and DNA hypomethylation, the death of the mouse Pm299L HCC cell line was preceded by a significant induction of stress-related genes (*ATF4* and *CHOP*), accumulation of NMD targets (*ATF3* and *SRSF3-ISO2*), and activation of the innate immune response (*IFN*β and *CXCL10*) (**Figure S1C**).

**Figure 1.**
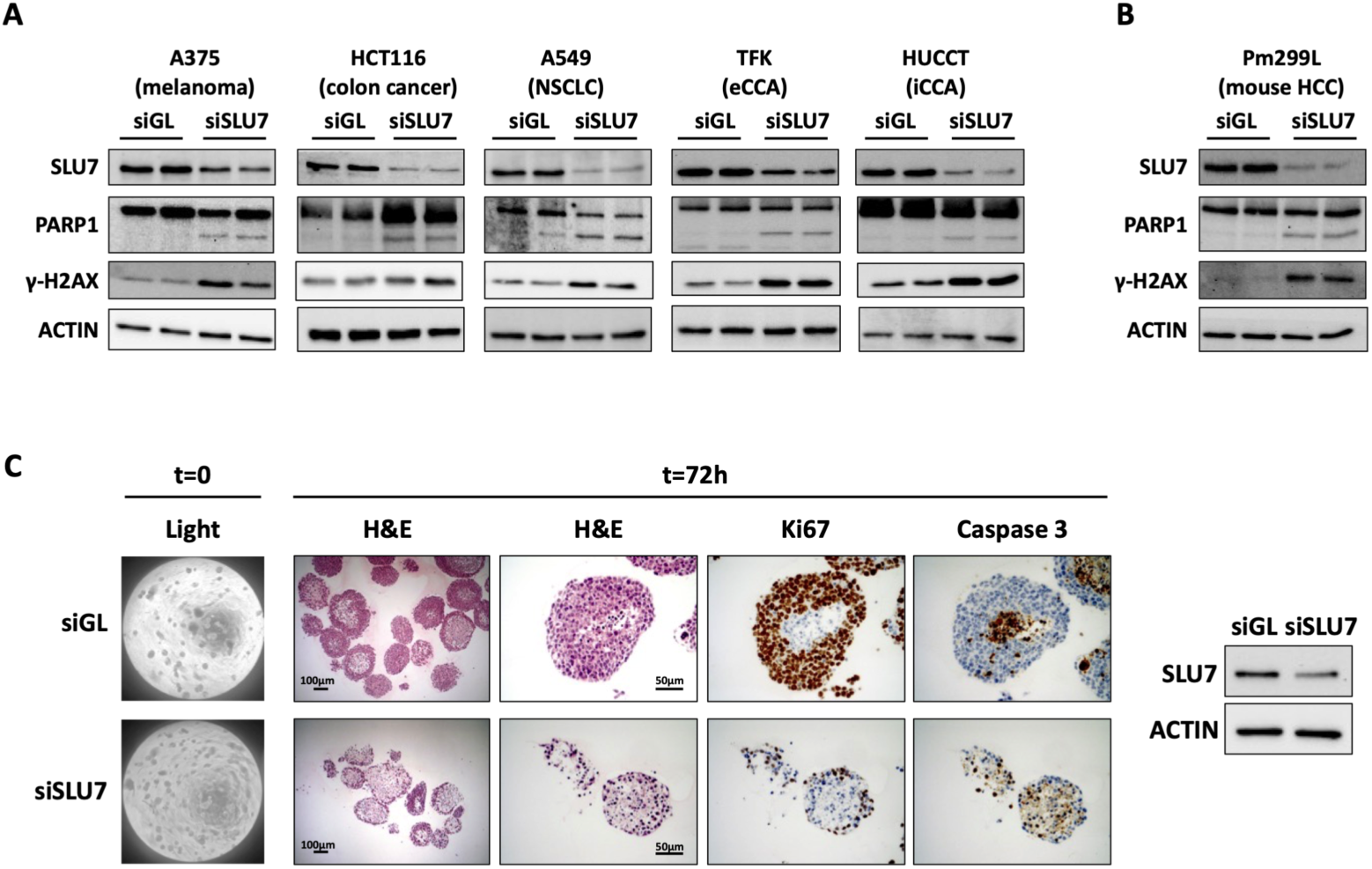
SLU7 silencing reduces human and mouse cancer cells viability. **A.** Silencing of SLU7 for 72 hours upon siSLU7 transfection led to increase PARP1 cleavage and γ-H2AX induction compared to control siRNA (siGL) transfection in human melanoma (A375), colon cancer (HCT116), NSCLC (A549), extrahepatic CCA (eCCA) (TFK-1), and iCCA (HUCCT-1) cells. The Western blot of Actin is shown as loading control. **B.** The same results were observed in the mouse HCC cell line Pm299L. **C.** Organoids were generated from a liver orthotopic tumor obtained in C57BL/6J mice upon implantation of a piece of a subcutaneous tumor induced with Pm299L HCC cells. Once organoids were established, SLU7 expression was silenced upon siRNA transfection and 72 hours after organoids were collected to analyze SLU7 expression by Western blot (right panel) and to be fixed and paraffin embedded for hematoxylin & eosin staining (H&E) and Ki67 and Caspase 3 immunohistochemistry.

To confirm this effect under more physiological conditions, we generated organoids from a liver orthotopic tumor obtained in C57BL/6J mice upon implantation of a piece of a subcutaneously induced Pm299L HCC cell line tumor. Once organoids were established (**Figure 1C**; t=0), SLU7 expression was silenced and 72 hours after siRNAs transfection organoids were collected to analyze SLU7 expression by Western blot and were fixed and paraffin embedded for further analyses.

As shown in **Figure 1C**, the H&E staining demonstrates the disruption of the organoids upon SLU7 silencing, which was accompanied by a significant reduction in the expression of Ki67 and the loss of the central distribution of caspase 3 expression. We then decided to test whether SLU7 silencing could be a good anti-tumoral strategy.

### 2. Intratumoral inducible SLU7 silencing reduces heterotopic syngeneic tumoral growth

To demonstrate that intratumoral SLU7 inhibition could be a potential anti-tumoral strategy, we first generated mouse HCC Pm299L cells expressing SLU7 specific shRNAs in a doxycycline (DOX) inducible manner (shSLU7i) or a non-targeting negative control shRNA (shNTCi), by lentivirus infection (**Figure S2A**).

After selecting and testing several clones, we confirmed that DOX treatment led to SLU7 expression inhibition accompanied by PARP1 cleavage, DNA damage and cell death in Pm299L-shSLU7i HCC cells but not in Pm299L-shNTCi HCC control cells (**Figure 2A and B**). To evaluate this effect *in vivo*, Pm299L-shSLU7i HCC cells were subcutaneously injected into C57BL/6J mice. Once the tumors were established, mice were divided into two groups for gavage administration of either PBS or DOX (**Figure 2C**). As shown in **Figure 2D**, upon daily administration for 9 days, tumor volume was significantly reduced in mice receiving DOX.

**Figure 2.**
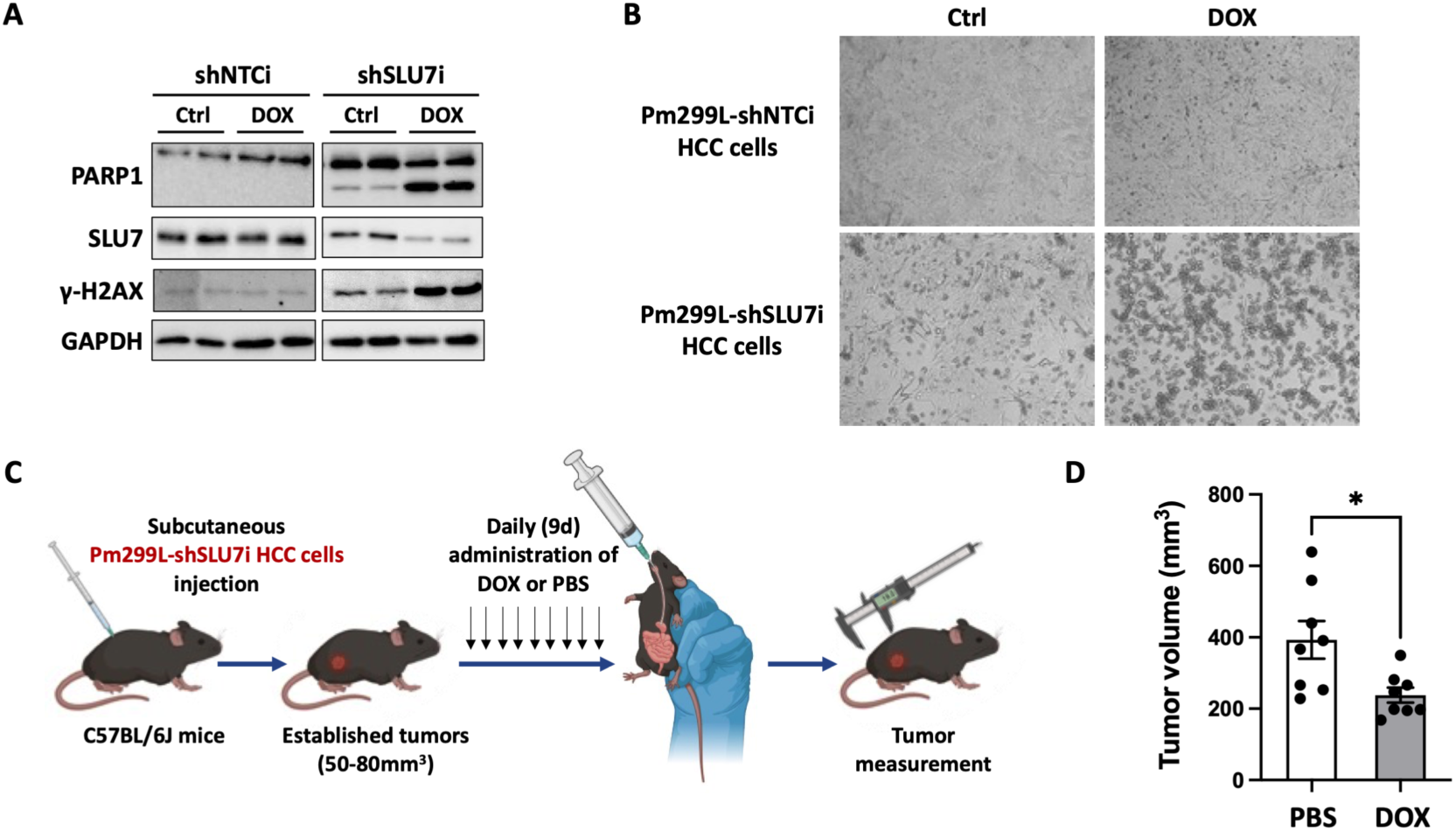
Intratumoral inducible SLU7 silencing inhibits the growth of subcutaneous syngeneic tumors. **A.** Western blot to study the effect of doxycycline (DOX) treatment on cell lines expressing an inducible control shRNA (shNTCi) or a SLU7 specific inducible shRNA (shSLU7i) on the cleavage of PARP1 and the induction of γ-H2AX. The expression of SLU7 and GAPDH are included as silencing and load controls respectively. **B.** Light microscopy images of cells analyzed in A. **C.** Treatment schedule to induce subcutaneous tumors in C57BL/6J mice with Pm299L-shNTCi HCC cells and to study the effect on the growth of established tumors of SLU7 silencing upon oral administration of doxycycline (DOX) to mice. **D.** The downregulation of SLU7 expression in established tumors upon DOX administration significantly reduced tumor volume.

### 3. Intratumoral inducible *SLU7* edition reduces heterotopic syngeneic tumor growth

We used CRISPR/Cas9 technology to study the effect of intratumoral SLU7 gene editing on tumor growth. To this end, we first generated stable clones of Pm299L HCC cells expressing DOX-inducible Cas9 (iCas9) nuclease. From these, we selected one clone showing high inducible Cas9 expression upon DOX treatment and undetectable expression levels under unstimulated conditions. This clone was then used to generate and select stable polyclonal cell populations constitutively expressing either a sgRNA directed to SLU7 gene (sgSLU7) or a non-targeting control sgRNA (sgNTC) (**Figure S2B**). As shown in **Figure 3A**, treatment with DOX of sgNTC cells had no significant effect on SLU7 expression, PARP1 cleavage, DNA damage and cell viability (**Figure 3B**). In contrast, DOX treatment of cells expressing the SLU7 sgRNA resulted in SLU7 expression inhibition, PARP1 cleavage, DNA damage and cell death (**Figure 3A-B**). As expected, these effects were accompanied by a significant edition of SLU7 gene in Pm299L-iCas9-sgSLU7 cells which was not observed in Pm299L-iCas9-sgNTC control cells upon DOX treatment (**Figure 3C**).

**Figure 3.**
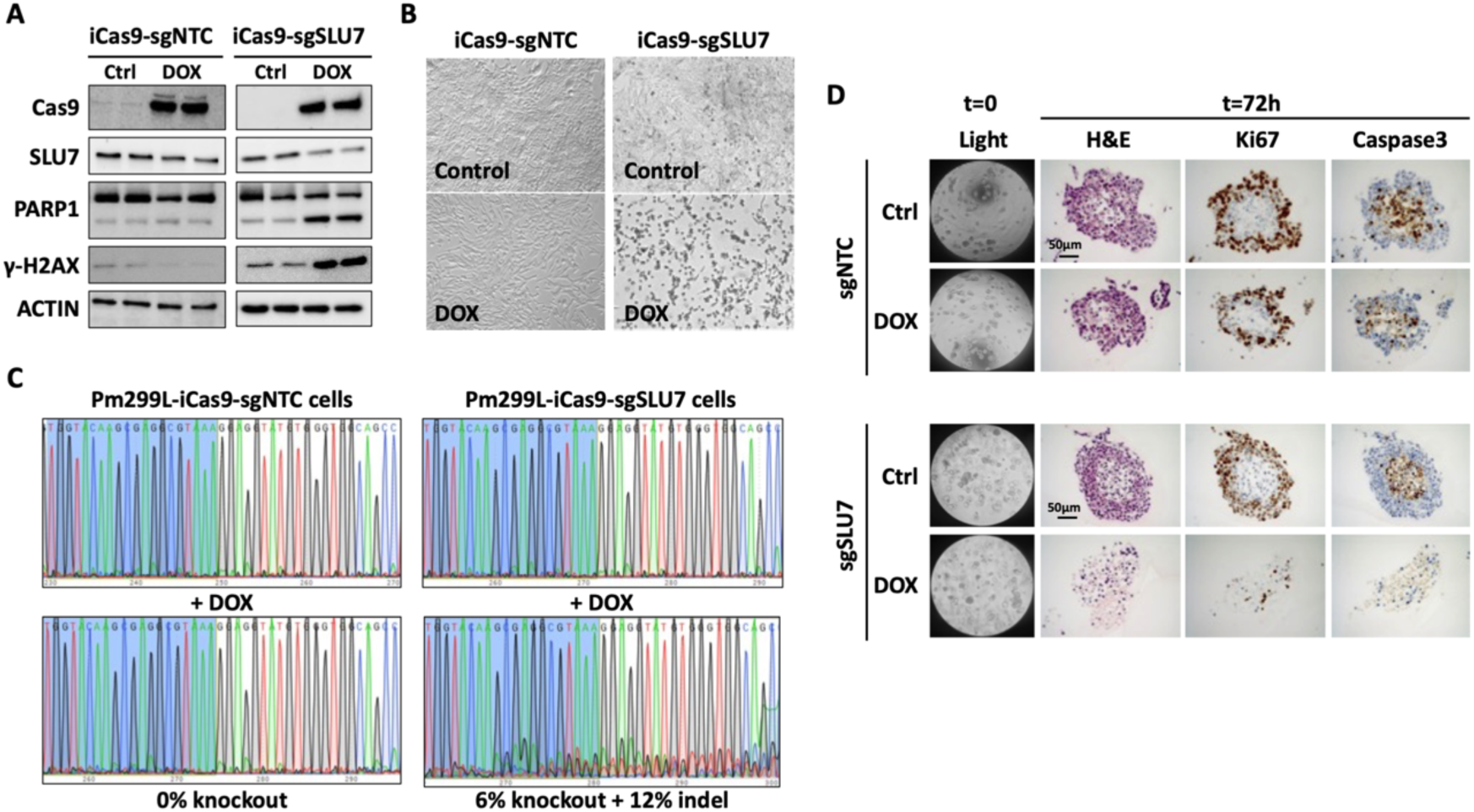
Inducible SLU7 gene edition induces the death of Pm299L HCC cells. **A.** Western blot of Cas9, SLU7, PARP1, γ-H2AX and Actin in Pm299L-iCas9-sgNTC control cells and Pm299L-iCas9-sgSLU7 in basal conditions and 72 hours after doxycycline (DOX) addition. The induction of Cas9 upon DOX treatment in Pm299L-iCas9-sgSLU7 cells is parallel to the silencing of SLU7, the cleavage of PARP1 and the induction of γ-H2AX. **B.** Viability of the cells analyzed in A. **C.** Sanger sequencing of SLU7 gene in the DNA obtained from cells analyzed in A, showing the SLU7 edition in Pm299L-iCas9-sgSLU7 cells upon DOX treatment. **D.** Organoids derived from liver tumors formed after intrahepatic implantation of a tissue fragment from a subcutaneous tumor previously induced with Pm299L-iCas9-sgSLU7 (sgSLU7) or Pm299L-iCas9-sgNTC (sgNTC) HCC cells, were cultured and analyzed after fixation and paraffin embedding. Hematoxylin & eosin (H&E) staining and immunohistochemistry detection of Ki67 and caspase 3 was performed in control conditions and 72 hours after treatment with DOX.

Before evaluating the impact of intratumoral SLU7 editing on tumor growth, we decided to assess the effect of SLU7 editing on tumor-derived organoids. These organoids were generated from liver tumors formed after intrahepatic implantation of a tissue fragment from a subcutaneous tumor previously induced with Pm299L-iCas9-sgSLU7 or Pm299L-iCas9-sgNTC HCC cells. As observed in **Figures 3D and S3**, a 72 hours DOX treatment has not a significant effect on the organoids derived from Pm299L-iCas9-sgNTC tumors. In contrast, DOX administration to organoids derived from Pm299L-iCas9-sgSLU7 tumors caused marked disorganization together with a significant reduction in proliferating and viable cells (**Figures 3D and S3**).

We then went on to test the *in vivo* antitumoral effect of SLU7 edition in established Pm299L-iCas9-sgSLU7 subcutaneous tumors. As shown in **Figure 4A**, DOX administration has no significant effect on the growth of tumors established by subcutaneous Pm299L-iCas9-sgNTC HCC cells injection, in which, as expected, no SLU7 gene editing was observed (**Figure 4B**). However, DOX administration significantly prevented the growth of Pm299L-iCas9-sgSLU7 subcutaneous tumors (**Figure 4C**), in parallel to SLU7 gene editing (**Figure 4D**).

**Figure 4.**
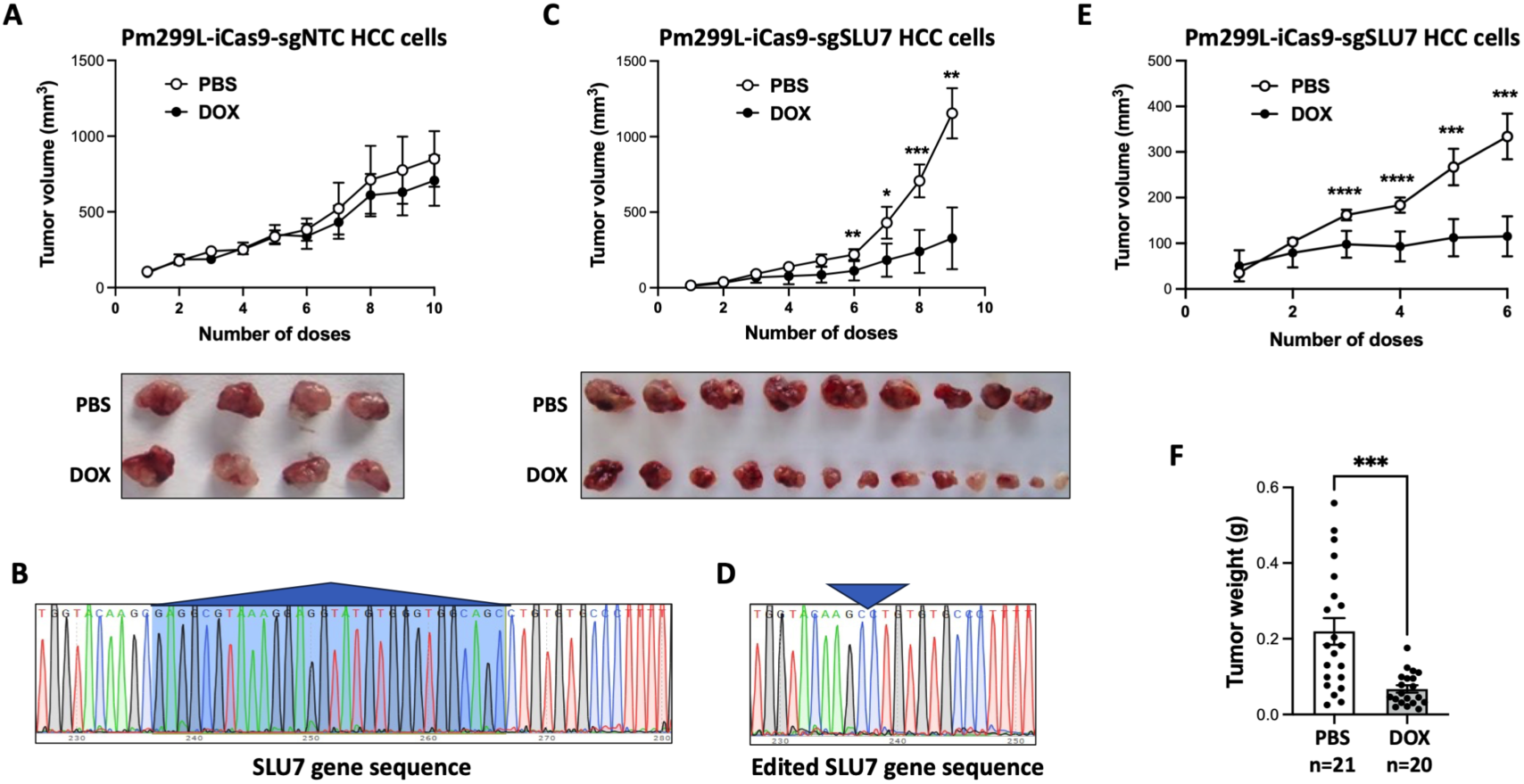
Intratumor SLU7 edition has antitumoral effect in established Pm299L-iCas9-sgSLU7 subcutaneous tumors. **A.** The growth of subcutaneous tumors induced in C57BL/6J mice with Pm299L-iCas9-sgNTC HCC cells was similar in mice orally administered with PBS or doxycycline (DOX). **B.** DOX treatment had no effect on SLU7 gene edition in tumors induced with Pm299L-iCas9-sgNTC HCC cells. **C.** The growth of subcutaneous tumors induced in C57BL/6J mice with Pm299L-iCas9-sgSLU7 HCC cells was significantly reduced in mice orally administered DOX in comparison with control PBS. **D.** DOX treatment resulted in SLU7 gene edition in tumors induced with Pm299L-iCas9-sgSLU7 HCC cells. Arrow head indicates the point where the normal SLU7 gene sequence highlighted in blue in B is deleted. **E.** The growth of subcutaneous tumors induced in athymic Balb/c nude mice with Pm299L-iCas9-sgSLU7 HCC cells was significantly reduced in mice orally administered DOX in comparison with control PBS. **F.** The weight of subcutaneous tumors induced in athymic Balb/c nude mice with Pm299L-iCas9-sgSLU7 HCC cells was significantly reduced in mice orally administered DOX in comparison with control PBS.

In order to evaluate the contribution of the immune system to the antitumoral effect of SLU7 editing, we induced the formation of subcutaneous tumors with Pm299L-iCas9-sgSLU7 HCC cells in athymic Balb/c Nude mice. As shown in **Figure 4E-F**, similarly to what we observed in the immune competent mice, the administration of DOX significantly prevented tumor growth in those animals suggesting a marginal role of T cells in the antitumoral effect of SLU7 downregulation.

### 4. Intratumoral inducible SLU7 edition reduces orthotopic liver tumor growth

A subcutaneous tumor induced after injection of Pm299L-iCas9-sgSLU7 cells in C57BL/6J mice was collected and fragmented in small pieces to be used for orthotopic mouse liver implantation. At day 6 after liver implantation and upon ultrasound evaluation of tumor size mice were divided into two groups to receive DOX or PBS as control (**Figure 5A**). DOX administration for 9 days significantly reduced tumor growth (**Figure 5B-C**) which also resulted in a significantly lower liver to body weight ratio (*i.e*. tumor burden) (**Figure 5C**).

**Figure 5.**
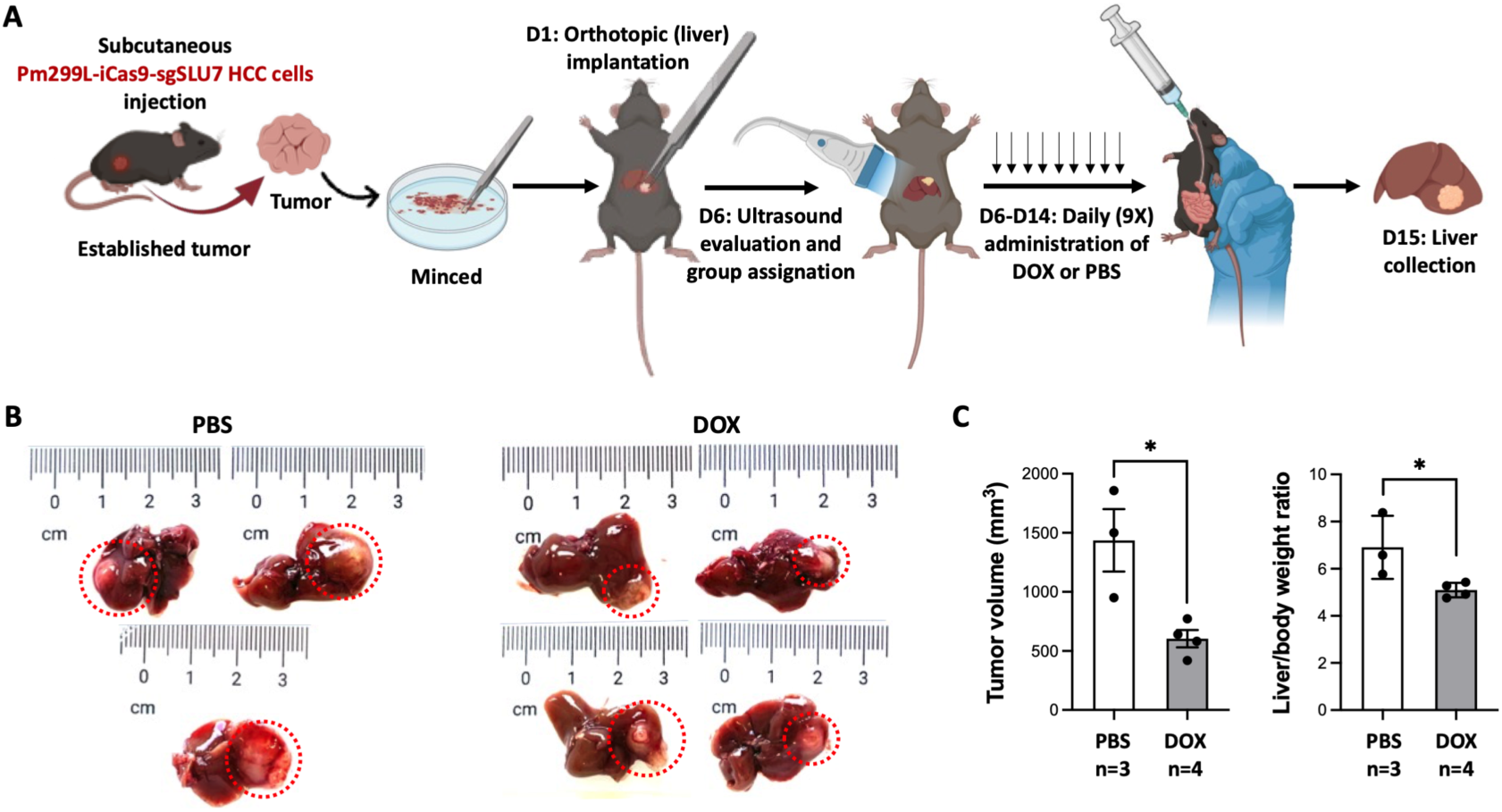
Antitumor effect of intratumoral SLU7 edition in orthotopic mouse HCC tumors. **A.** Schedule of tumor induction and treatment. A subcutaneous tumor was induced in C57BL/6J mice with Pm299L-iCas9-sgSLU7 HCC cells. The tumor was isolated and minced and small fragments were implanted in the left lobe of the liver of C57BL/6J mice. Tumor growth was followed by ultrasound examination and mice were divided in two groups to receive by gavage nine daily doses of PBS or doxycycline (DOX). At day 15 mice were sacrificed and tumors were measured and weighted. **B.** Liver tumors of mice treated with PBS or DOX as described in A. **C.** Tumor volume and the ratio of liver to body weight were significantly reduced in mice treated with DOX.

### 5. The systemic administration of a siSLU7 aptamer chimera (APTASLU) reduces orthotopic HCC tumor growth

At this point we designed a chimeric aptamer based on the DNA G4 quadruplex structure AS1411 directed to nucleolin. This is the first aptamer used in cancer clinical trials that demonstrated to be safe and well tolerated by patients^12^ that can be used to deliver siRNAs to tumors^5^, and which we conjugated to a SLU7 specific siRNA (APTASLU). To generate the APTASLU chimera the 3’ end of AS1411 DNA was elongated with the RNA sequence containing 2’-fluoro-modified pyrimidines of the passenger strand specific for mouse and human SLU7 and the aptamer was further hybridized with the guide strand with no modifications. A control chimera aptamer was also generated upon conjugation of AS1411 with siGL (APTAsiGL).

Although it has been shown that AS1411 binds to a plethora of human and mouse solid tumor cell lines expressing nucleolin at the surface, we first tested its ability to bind to Pm299L cells. As shown in **Figure S4A**, we labelled the AS1411 aptamer with AlexaFluor 647 and demonstrated its binding to Pm299L cells by flow cytometry. Then to confirm SLU7 silencing by APTASLU, we transfected Pm299L cells with lipofectamine or added directly the aptamers into the culture medium for free uptake. In both cases APTASLU downregulated SLU7 expression (**Figure S4B**).

We then tested the effect of APTASLU *in vivo*. We generated orthotopic HCC tumors as previously described (**Figure 6A**) and once the tumors were established and measured by ultrasound examination, animals were divided into two groups to receive retroorbital injection of 5 doses of APTAsiGL or APTASLU every two days. Eighteen days after liver orthotopic tumor implantation and after 5 systemic administrations of the aptamer chimeras’ mice were sacrificed and livers were collected. As shown in **Figure 6 B-C**, the liver tumors in mice receiving systemic APTASLU were significantly smaller than tumors of mice receiving APTAsiGL. Interestingly, when we analyzed the immune infiltrate in the excised tumors by immunohistochemistry, we found a significantly higher infiltration of CD3+ lymphocytes in APTASLU treated mice (**Figures 6D and S5)**, suggesting that SLU7 knockdown promotes tumor inflammation.

**Figure 6.**
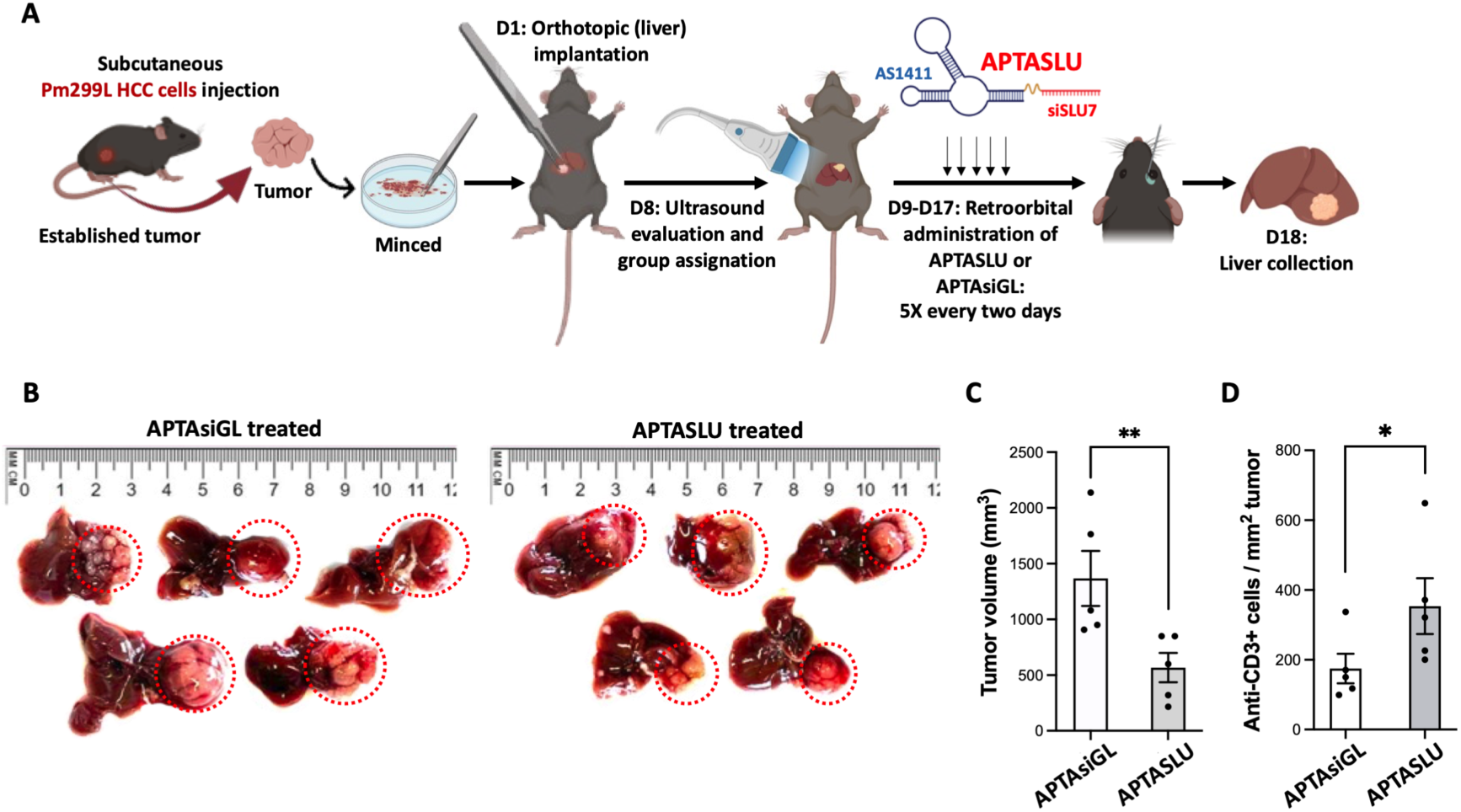
The systemic administration of APTASLU has antitumoral effect on orthotopic mouse HCC tumors. **A.** Schedule of tumor induction and treatment. A subcutaneous tumor was induced in C57BL/6J mice with Pm299L HCC cells. The tumor was isolated and minced and small fragments were implanted in the left lobe of the liver of C57BL/6J mice. Tumor growth was followed by ultrasound examination and mice were divided in two groups to receive by retroorbital injection APTASLU or APTAsiGL as control. At day 18 mice were sacrificed and tumors were measured. **B.** Liver tumors in mice treated with APTAsiGL or APTASLU as described in A. **C.** The volume of liver tumors was significantly reduced in mice treated systemically with APTASLU. **D.** Quantification of CD3+ cells identified in the tumors by immunohistochemistry.

### 6. Systemic APTASLU potentiates the antitumoral effect of anti-PD1 immunotherapy in a liver orthotopic tumor model

As in many other tumors immunotherapy with ICIs has significantly improved the prognosis of patients with advanced HCC, however, still a large proportion of patients do not respond to these therapies. Therefore, we decided to test whether APTASLU treatment could potentiate the antitumoral effect of anti-PD1 therapy. Once again, we used the orthotopic liver tumor model to compare the therapeutic effect of intraperitoneal administration of anti-PD1 alone or in combination with APTASLU administered retro-orbitally on alternate days (**Figure 7A**). Our results indicate that APTASLU may act as an immune enhancing strategy. Interestingly, although differences in tumor size were not statistically significant, only 1 out of 5 mice treated with the combination had a large tumor, 3 mice had very small tumors and no tumor was detected in another animal. On the contrary, in the group treated with anti-PD1 alone, all mice developed detectable tumors and 4 out of 5 mice exhibited very large tumors (**Figure 7B**).

**Figure 7.**
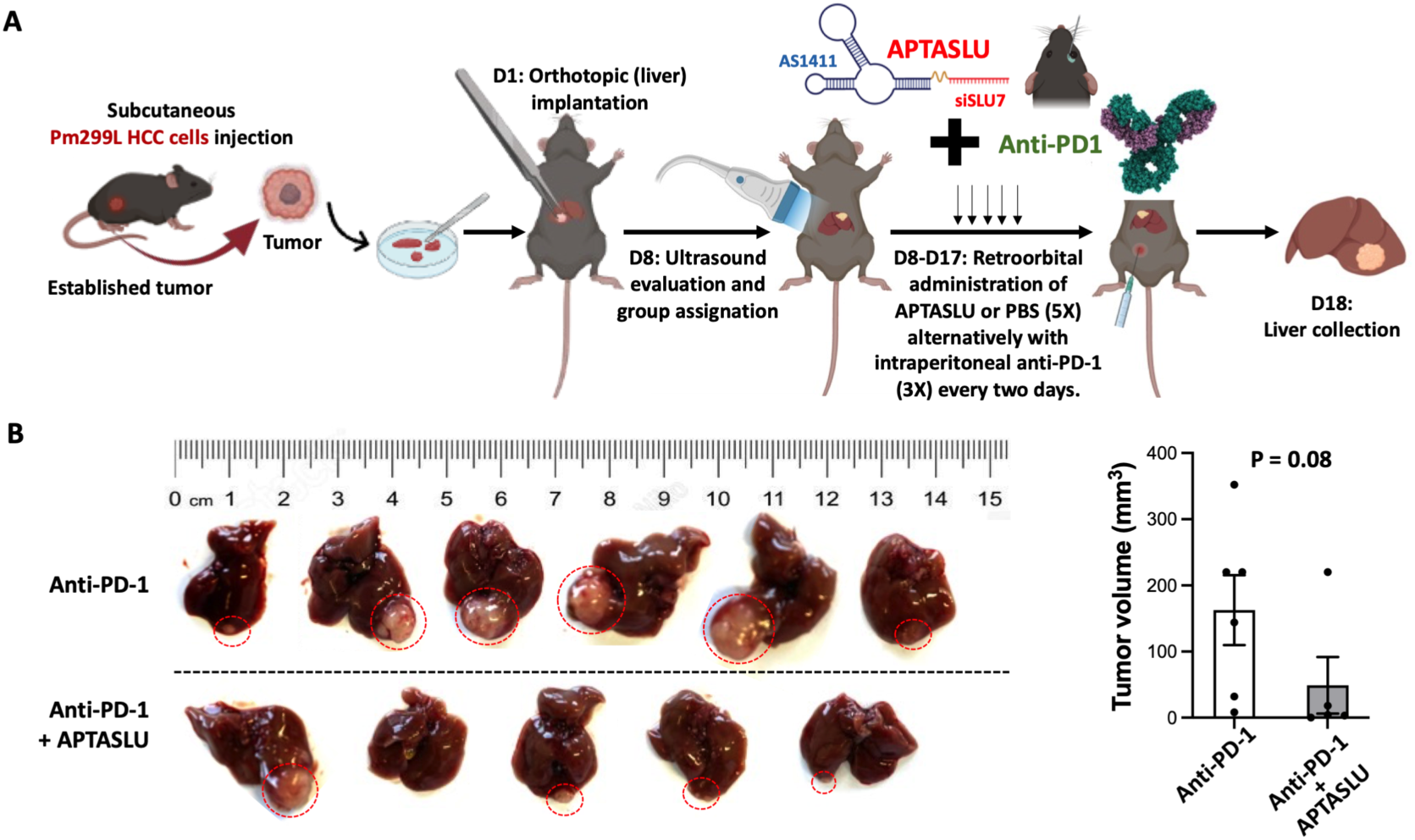
APTASLU potentiates the antitumoral effect of PD1 immune checkpoint blockade on orthotopic mouse HCC tumors. **A.** Schedule of tumor induction and treatment. A subcutaneous tumor was induced in C57BL/6J mice with Pm299L HCC cells. The tumor was isolated and minced and small fragments were implanted in the left lobe of the liver of C57BL/6J mice. Tumor growth was followed by ultrasound examination and mice were divided in two groups to receive intraperitoneally anti-PD-1 and by retroorbital injection APTASLU or PBS as control. At day 18 mice were sacrificed and tumors were measured. **B.** Liver tumors in mice treated with Anti-PD-1 or Anti-PD-1 plus APTASLU as described in A. **C.** Volume of liver tumors induced in C57BL/6J and treated as described in A.

### 7. Intratumoral injection of siSLU7 significantly reduces the growth of subcutaneous xenografts of human HCC, iCCA and colon cancer tumors

To further confirm the antitumoral potential of SLU7 silencing in human tumors we induced different subcutaneous xenograft tumors with human HCC (PLC/PRF/5), intrahepatic CCA (iCCA) (HUCCT-1) and colon cancer (HCT116) cells (**Figure 8A**). Once tumors were established, we intratumorally injected siSLU7 oligos (twice every 2 days) incorporated into magnetic nanoparticles, holding a magnet at the injection site in order to increase the retention of the nanoparticles in the tumor area (**Figure 8A**). In agreement with all our previous observations, the administration of just two doses of siSLU7 significantly reduced the growth of the tumors generated by injection of PLC/PRF/5 HCC cells (**Figure 8B**), HUCCT-1 iCCA cells (**Figure 8C**) and HCT116 colon cancer cells (**Figure 8D**).

**Figure 8.**
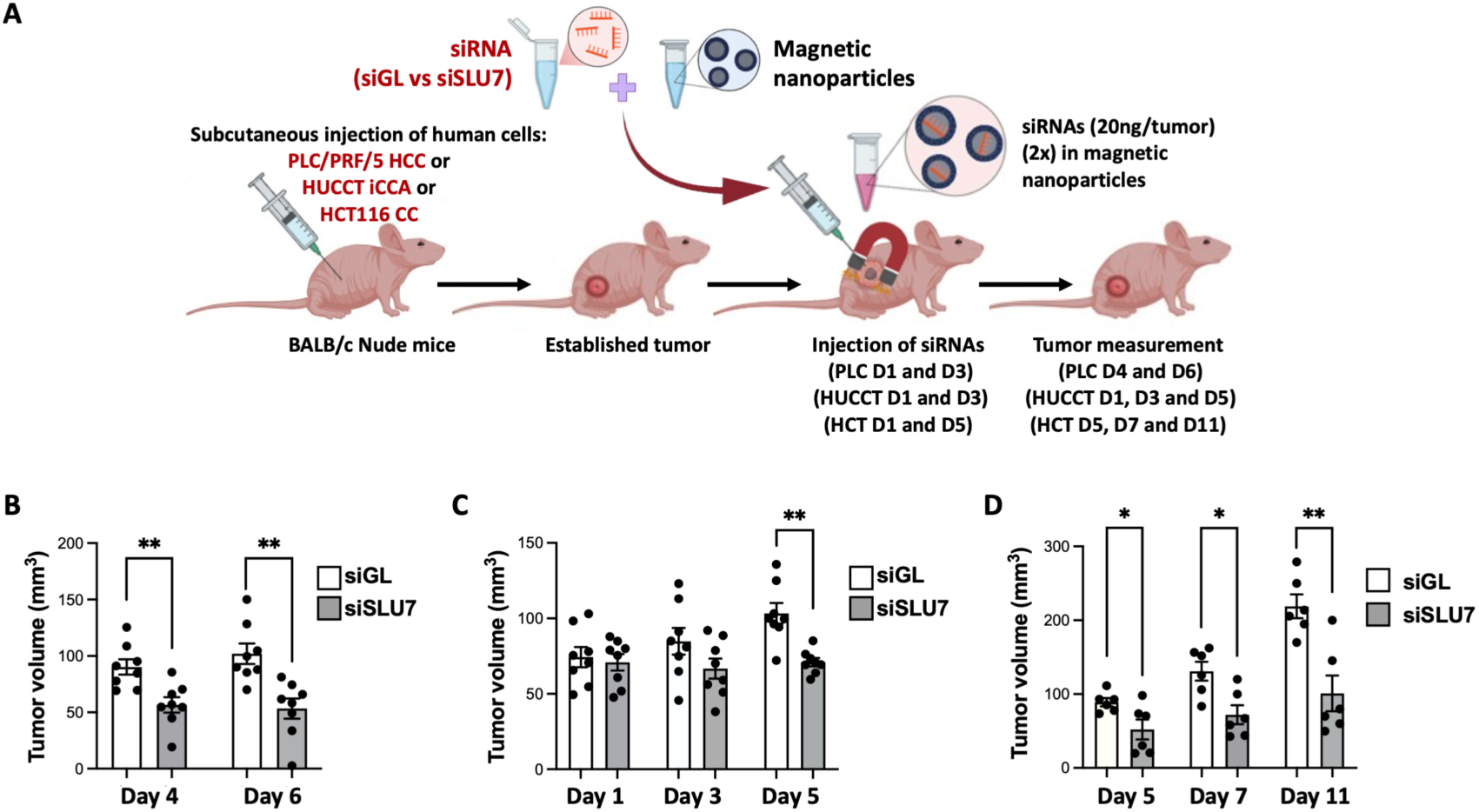
Intratumoral siSLU7 administration reduces the growth of subcutaneous human xenograft tumors. **A.** Schedule of tumor induction and treatment. Subcutaneous tumors were induced in Balb/c nude mice upon injection of human cancer cells PLC/PRF/5 (HCC), HUCCT-1 (iCCA) or HCT116 (colon cancer). Once tumors were established siSLU7 or siGL as negative control conjugated in magnetic particles were injected intratumorally in the presence of magnets. **B-D.** The volume of tumors induced with PLC7PRF/5 HCC (**B**), HUCCT-1 iCCA (**C**) or HCT116 colon cancer (**D**) human cells was significantly inhibited upon siSLU7 treatment.

## DISCUSSION

We have identified SLU7 as an essential, non-redundant survival factor not only for HCC cells, but also for other mouse and human cancer cells of diverse origins. This is consistent with evidence that multiple molecular cancer subtypes depend on proper splicing function for survival^13^. Notably, SLU7 knockdown does not compromise the viability of normal hepatocytes^6^. Mechanistically, SLU7 silencing in transformed cells induces apoptosis, which is preceded by autophagy, oxidative stress, the accumulation of RNA-DNA hybrids (R-loops), and DNA damage^7,8,14^. R-loops, in particular, can disrupt DNA replication and promote genomic instability and replication stress, features that represent exploitable therapeutic vulnerabilities in cancer cells^15,16^.

Additionally, SLU7 knockdown leads to aberrant splicing isoforms and NMD mRNA targets accumulation due to the inhibition of splicing and NMD^10^. Again, targeting both splicing and the NMD RNA surveillance pathway are emerging strategies to expand the tumor antigen repertoire^17,18^, which could be leveraged to enhance the efficacy of cancer immunotherapy.

According to all these data, we present clear *in vivo* evidence of the anti-tumoral effect of SLU7 knockdown alone or in combination with immune checkpoint inhibitors, across various mouse models, including subcutaneous and orthotopic liver models in syngeneic immune-competent mice, and subcutaneous xenograft of different human tumors.

As proof of concept, we used inducible CRISPR/Cas9 approaches to demonstrate that SLU7 editing in tumor cells of established tumors is sufficient to inhibit subcutaneous cancer growth, both in immune competent and immune compromised mice, as well as orthotopic liver tumor growth.

Furthermore, we explored different non-coding RNA-based therapeutic strategies directed to modify the expression of selected genes, with potential utility to treat cancer^19^. Among them, siRNAs can trigger sequence-specific cleavage of mRNA molecules, however, their efficient delivery to tumor cells is essential to reduce side effects and to enhance efficacy. One delivery method involves aptamer-siRNA chimeras^20^. Aptamers are short single-stranded DNA (ssDNA) or RNA molecules that due to their 3D conformations are able to bind a wide range of molecular targets with high affinity^20,21^. AS1411, a well characterized 26-mer DNA aptamer with G-quadruplex structure that binds surface-expressed nucleolin in many transformed cells, has been validated in multiple clinical trials as both a targeting ligand and therapeutic agent^22^. Here we synthesized APTASLU, a chimeric oligonucleotide comprising AS1411 and an antisense SLU7 siRNA strand hybridized with its passenger strand. APTASLU effectively silenced SLU7 expression and importantly, when systemically administered, significantly reduced orthotopic liver tumor growth increasing tumor inflammation. Accordingly, APTASLU enhanced the anti-tumor efficacy of anti-PD-1 treatment in this model.

As an alternative delivery approach, we employed nanoparticles to deliver siSLU7 oligos to cancer cells. To maximize silencing efficiency and minimize systemic dissemination and toxicity, we injected siSLU7-loaded magnetic nanoparticles into magnetically targeted subcutaneous tumors. Remarkably, just two doses were sufficient to significantly reduce tumor growth in subcutaneous models of human HCC, iCCA and colon carcinoma.

In summary, our findings show that *in vivo* targeting of SLU7 is both feasible and effective in eliciting anti-tumor responses. These effects can be achieved through various molecular strategies, siRNA sequences, and delivery methods and could be combined to improve the efficacy of immunotherapy. Further research is needed to better understand the underlying molecular mechanisms.

## Material and Methods

### Cell lines

Human cell lines PLC/PRF/5 (HCC), A375 (melanoma), HCT116 (colon cancer), A549 (NSCLC), HUCCT-1 (iCCA), TFK-1 (eCCA), mouse cell lines Hepa 1.6 (HCC), B16/F10 (melanoma), and 4T1 (breast cancer) were obtained from the American Type Culture Collection (ATCC). All cell lines were authenticated by short tandem repeat (STR) profiling and routinely tested for mycoplasma contamination using the *MycoAlert* Mycoplasma Detection Kit (Lonza, #LT07-318). The mouse HCC cell line Pm299L was established previously by Dr. Amaia Lujambio (Mount Sinai, New York, USA). PLC/PRF/5, A375, TFK-1, HUCCT-1, B16/F10 and Hepa 1.6 cells were cultured in Advanced DMEM (Gibco-Life Technology, Madrid, Spain) supplemented with 10% fetal bovine serum (FBS), glutamine, and antibiotics. Pm299L and 4T1 cells were maintained in DMEM (Gibco-Life Technology) supplemented with 10% FBS, glutamine, and antibiotics. HCT116 cells were grown in McCoy’s 5A medium (Gibco-Life Technology) supplemented with 10% FBS, glutamine, and antibiotics. A549 cells were cultured in RPMI (Gibco-Life Technology) supplemented with 10% FBS, glutamine, and antibiotics. All cells were maintained in a humidified incubator at 37°C with 5% CO₂.

### Generation of lentiviral particles

The different lentiviral particles were produced by transfection of HEK293T cells with the corresponding plasmids as previously described^23^. After filtered through a 0.2 μm cellulose-free filter (Sartorius, Göttingen, Germany) particles were directly applied to target cells at a multiplicity of infection (MOI) <1.

### Generation of inducible SLU7 knockdown cell lines

To generate a CRISPR-Cas9 inducible knockdown model, Pm299L cells were first transduced with lentiviral particles encoding a doxycycline (DOX)-inducible Cas9 endonuclease (Lenti-iCas9-neo, Addgene plasmid #85400). After selection with 800 μg/mL neomycin (InvivoGen, San Diego, CA, USA #108321-42-2), GFP-positive cells were sorted following DOX treatment using a MoFlo Astrios EQ sorter (Beckman Coulter), seeded individually into 96-well plates, expanded, and evaluated for DOX-induced GFP expression by flow cytometry. A single clone showing robust inducible Cas9 expression referred as Pm299L-iCas9 was selected for subsequent transduction with sgRNA-expressing lentiviral vectors.

Single-guide RNAs (sgRNAs) targeting the *SLU7* gene at exon 3, were designed using the Integrated DNA Technologies (IDT; https://eu.idtdna.com/page) online tool and cloned into the Lenti-sgRNA-puro vector (Addgene plasmid #52963). A non-targeting sgRNA (sgNTC) was also cloned as control. Pm299L-iCas9 cells were then infected with lentiviruses encoding either sgNTC or sgSLU7 and maintained under dual selection with 800 μg/mL neomycin and 6 μg/mL puromycin (InvivoGen, #58-58-2). Resulting clone pools (either Pm299L-iCas9-sgSLU7 or Pm299L-iCas9-sgNTC) were expanded and characterized for SLU7 gene editing by Sanger sequencing and for SLU7 protein depletion by Western blot after DOX treatment.

To generate the inducible shRNA knockdown model (Pm299L-shSLU7i), short hairpin RNAs (shRNAs) targeting SLU7 were designed using the Broad Institute’s GPP Web Portal (https://portals.broadinstitute.org/gpp/public/) and cloned into the Tet-pLKO.1-puro vector (Addgene plasmid #21915). A non-targeting shRNA (shNTC) was used as control. Pm299L cells were infected with the corresponding lentiviral particles and selected using 6 μg/mL puromycin. Resulting shNTCi and shSLU7i pools of clones were characterized by Western blotting to confirm SLU7 depletion after DOX treatment. To reduce potential interference with possible traces of tetracycline in standard serum, all cell lines in this section, including HEK293T and Pm299L-derived models, were cultured with 10% FBS FetalClone (Cytiva, Marlborough, MA, USA; #SH30109.03).

### Generation and characterization of aptamer AS1411-siRNA conjugates

We first validated the binding of AS1411 aptamer to Pm299L cells by flow cytometry using a AS1411 conjugated to AlexaFluor 647. Cells were incubated with 40 pmol of AlexaFluor-AS1411 DNA for 30 min at 37°C in PBS, washed twice, and analyzed in the CytoFLEX LS flow cytometer (Beckman Coulter). Data were processed using FlowJo v10 (FlowJo LLC).

Characterization of AS1411-siRNAs conjugates: AS1411 ssDNA aptamer (dGdGdTdGdGdTdGdGdTdGdGdTdTdGdTdGdGdTdGdGdTdGdGdTdGdG), extended at the 3ʹ end with exon 9 SLU7 siRNA (rG/i2FC/rArA/i2FU/rGrArGrArGrArGrArA/i2FU//i2FC/ /i2FC//i2FU//32FU/) (APTA-SLU) or a control siRNA (i2FC/rG/i2FU/rA/i2FC/rG/i2FC/ rGrGrArA/i2FU/rA/i2FC//i2FU//i2FU//i2FC/rGrA) (APTA-siGL) were purchased from Integrated DNA Technologies (IDT) with the indicated 5-fluorouracil modifications in pyrimidines. Antisense (guide strand) siRNAs (siSLU7 rArArGrGrArUrUrCrUrCrUrC rUrCrArUrUrGrC and siGL rUrCrGrArArGrUrArUrUrCrCrGrCrGrUrArCrG) were hybridized in annealing buffer (NaCl 0.15 M; EDTA 0.01 M; Tris-Cl pH 8.8; adjusted to final pH = 7.5) in a thermocycler starting at 65°C and allowed to cool to 37°C. Correct hybridization of APTA-SLU or APTA-siGL was checked in a 15% acrylamide SDS-PAGE. Expected molecular weight was assessed with RNA Marker Low (Abnova). Next, we checked in vitro SLU7 silencing by APTASLU in Pm299L cells after free aptamer uptake or lipofectamine 2000 (Invitrogen)-mediated transfection of 0.5 µM APTAsiGL or APTASLU, following the manufacturer’s instructions. For free uptake assays, cells were incubated with 0.5 µM APTAsiGL or APTASLU diluted in 500 μL of Opti-MEM (Gibco) for 2 hours. Then, 1 mL of DMEM (Gibco-Life Technology) supplemented with 10% FBS, glutamine, and antibiotics was added for 2 hours. The same treatment with APTAsiGL and APTASLU was repeated 6 hours after medium addition. Following a final 2-hour incubation with the aptamer–siRNA conjugates, 2 mL of complete DMEM was added, and cells were maintained for 48 hours before downstream analyses.

### Mouse tumor models

All animal used received humane care according to the criteria outlined in the ‘Guide for the care and Use of Laboratory Animals’ prepared by the National Academy of Sciences and published by the National Institutes of Health (NIH publication 86-23 revised 1985). Protocols were also approved and performed according to the guidelines of the Ethics Committee for Animal Testing of the University of Navarra (CEEA-130/19) and Navarra Government (GN 164/2020).

Subcutaneous tumor models were established in 6–8-week-old male mice. For syngeneic models, 7 x 10^5^ Pm299L-shSLU7i, Pm299L-iCas9-sgSLU7, or Pm299L-iCas9-sgNTC cells were implanted in the dorsal flank of C57BL/6J mice (Envigo, Huntingdon, UK). Once tumors reached an approximate volume of 50 mm³, mice were randomly divided into two groups to be treated daily by oral gavage with either doxycycline (DOX, 2 mg/mouse) or PBS as a control. The number of doses depended on the tumor type and it is indicated in the corresponding figure legend. In xenograft models, 10 x 10⁶ PLC/PRF/5, 5 x 10⁶ HUCCT-1, or 2 x 10⁶ HCT116 human cells were subcutaneously injected in the dorsal flank of BALB/c nude mice (Envigo). Once tumors reached an approximate volume of 50 mm³, tumors were locally treated twice with 20 µg of siSLU7-loaded into *In vivo SilenceMag* magnetic nanoparticles or siGL as a control (OzBiosciences, Marseille, France, #IV-SM31000), following the manufacturer’s instructions. The frequency of the treatment is indicated in the corresponding figure. Tumor volumes were measured everyday by caliper and mice were sacrificed under institutional approved protocols. Tumor volumes were calculated using the following formula: [length x (width)^2^)/2].

Orthotopic liver tumor models were generated as follows: first, subcutaneous syngeneic models were established by injecting Pm299L or Pm299L-iCas9-sgSLU7 cells into C57BL/6J mice (Envigo) as previously described. Once tumors were established, they were excised, and a 1 mm³ tumor fragment was implanted into the left liver lobe of recipient C57BL/6J mice (Envigo). Tumor growth was confirmed using VEVO3100 ultrasound imaging system (Fujifilm Visualsonics, Toronto, Canada).

For the Pm299L-iCas9-sgSLU7 syngeneic model, mice were randomized into two groups with similar tumor size and treated daily by oral gavage with either DOX at 2 mg/mouse or PBS as a control.

In experiments using Pm299L cells, mice received intravenously via retro-orbital injection SLU7 siRNA conjugated to the AS1411 aptamer (APTASLU7) or control APTAsiGL at a dose of 400 pmol per injection, administered five times every other day. When indicated, anti-PD1 treatment (BioXCell, New Hampshire, USA, BE0146), was administered at a dose of 80 µg per mouse intraperitoneally three times every two days.

### Mouse tumor-derived organoids

Organoids were derived from the orthotopic liver tumors generated with Pm299L, Pm299L-iCas9-sgSLU7 cells or control Pm299L-iCas9-sgNTC cells, as described above. After tumor excision, ∼50 mm³ fragments were dissected and enzymatically digested in collagenase/DNase solution (Roche, Basel, Switzerland, #11088866001 and #11284932001) for 30 minutes at 37°C. The resulting cell suspension was filtered through a 100 μm cell strainer (Corning, NY, USA #352360), and the enzymatic reaction was stopped by adding AutoMax buffer (prepared by supplementing 500 mL of PBS (Gibco) with 2 mL of fetal calf serum (Serana, Pessin, Germany, #00019560], 2.5 mL of 0.5 M EDTA, and 5 mL of penicillin-streptomycin). To lyse erythrocytes, cells were incubated with ACK lysis buffer (150 mM NH₄Cl, 10 mM KHCO₃, 0.1 mM EDTA) for 2 minutes at 37 ᵒC, the enzymatic reaction was stopped by adding cold medium. Cells (85,000 cells per drop) were then washed twice with Advanced DMEM/F12 medium, centrifuged at 300 × g for 5 minutes, resuspended in 30 μL of phenol red-free Matrigel (Corning, Arizona, USA, #356237) and cultured as described^24^.

For passaging, organoids were collected into 15 mL tubes and digested with 1 mL of TrypLE Express Enzyme 1X (Gibco, #12604-021) at 37°C for 5 minutes. Cells were then washed with medium and centrifuged at 300 × g for 5 minutes at 4°C. The pellet was resuspended in Matrigel and seeded as described above, using a 1:2 to 1:3 split ratio depending on the growth rate.

Organoids were characterized by brightfield microscopy using a Cell Observer Z1 (Carl Zeiss), immunohistochemistry, PCR and Western blot analysis. Previous to protein and DNA extraction, organoids were enzymatically dissociated with TrypLE Express Enzyme.

### siRNA transfections

All siRNAs were obtained from Sigma-Aldrich and used at a final concentration of 75 nM. Silencing efficiency was confirmed by Western blot. Sequences of SLU7 siRNAs, targeting exons 8 or 9/10, are available upon request. Transfections were performed using Lipofectamine RNAiMAX reagent (Invitrogen, Grand Island, NY, USA, #13778075) according to the manufacturer’s instructions.

For cell lines, transfections were carried out directly in adherent cultures following the standard RNAiMAX protocol.

For organoids, transfections were performed once they were fully established (5-7 days after passing). Briefly, the Matrigel drop containing the organoids was gently collected and transferred to a microcentrifuge tube containing PBS. The suspension was centrifuged at 200 x g for 1 minute at 4°C. After carefully removing the supernatant, organoids were washed once more with PBS. Following the second wash, PBS was completely and carefully removed, and organoids were incubated with the siRNA–Lipofectamine RNAiMAX transfection mixture in Eppendorf tubes placed on a rotator incubated at 32°C for 5 hours. After incubation, organoids were centrifuged at 200 x g for 1 minute and re-embedded in fresh phenol red-free Matrigel for culture.

### Immunohistochemical analysis of organoids

For histological processing, Matrigel drops containing organoids were carefully collected in PBS and washed twice by centrifugation at 200 x g for 1 minute at 4°C. Organoids were then fixed for 1h at room temperature in 4% paraformaldehyde (PFA; Panreac Applichem, #252931.1214) and embedded in 2% agarose prior to paraffin embedding. Paraffin blocks were sectioned at 3 μm thickness and processed for hematoxylin and eosin (H&E) staining, as well as immunohistochemistry for cleaved caspase-3 (Cell Signaling, 9661; 1:100) and Ki-67 (Cell Signaling, 12202; 1:500). In addition, immunohistochemistry for CD3 (Invitrogen, MA1-90582; 1:200) was performed on orthotopic liver tumors from mice; this staining, as well as all the organoid stainings, was carried out at the Morphology Core Facility at CIMA. Slides were scanned using an Aperio CS2 digital scanner (Leica Biosystems). Quantification of CD3 positive cells was performed using QuPath Software version 0.5.0 (University of Edinburgh) utilizing the “Positive cell detection” tool across all tumoral areas of the slides, at the magnifications indicated in the figures.

### Protein extraction and Western blot analysis

For protein extraction, cell lines and organoids were lysed in RIPA buffer (5 M NaCl, 1 M Tris, 0,5% Deoxycholate, 20% SDS, 1% Triton X-100 and a cocktail of phosphatases (1 mM sodium orthovanadate, 10 mM sodium fluoride, 100 mM β-glycerophosphate) and proteases inhibitors (Roche, Basel, Switzerland)) for 20 min at 4°C under constant rotation, sonicated and centrifuged at 12000 rpm for 20 min at 4°C.

All protein extracts were subjected to Western blot analysis as reported^8^. The antibodies used were: PARP1 (Cell Signaling, #46D11, 1/1000), ACTIN (Sigma-Aldrich, #A2066, 1/5000), γ-H2AX (Cell Signaling, #S139, 1/1000), GAPDH (Cell Signaling, #2118, 1/5000), SLU7 (Novus IBiologkalcS, #NBP2-20403, 1/1000) and Cas9 (Cell Signaling, #7A9-3A3, 1/1000).

### Total RNA isolation and PCRs

Total RNA from cell lines was extracted using the Maxwell RSC Instrument with simplyRNA tissue kit (Promega, Madison, WI, USA, AS1340). RNA samples were treated with DNAse to degrade all possible traces of contaminating genomic DNA (gDNA). Reverse transcription was performed as previously described^25^. Real-time PCRs were performed using an iCycler (Bio-Rad, Hercules, CA, USA) and the iQ SYBR Green Supermix (Bio-Rad) as previously described^26^. To monitor the specificity final PCR products were analyzed by melting curves and the amount of each transcript was expressed relative to the housekeeping gene RPLP0 as 2ΔCt, where ΔCt represents the difference in threshold cycle between the control and target genes, as described^27^. The sequence of primers used in the study will be provided upon request.

### Total DNA Isolation

Total DNA from cells, organoids and tumors was isolated using the Maxwell® RSC Cultured Cells DNA Purification Kit with a Maxwell® RSC 48 instrument (Promega, #AS1620). DNA of Pm299L-iCas9-sgSLU7 or control Pm299L-iCas9-sgNTC -derived subcutaneous tumors was extracted after 96h of DOX treatment from sorted (MoFlo Astrios EQ sorter, Beckman Coulter) GFP-positive cells. DNA purity and concentration were measured using a NanoDrop spectrophotometer (Thermo Fisher Scientific, Waltham, MA, USA).

### PCR and Sanger sequencing

PCRs of SLU7 were performed using 5 ng of genomic DNA, the Platinum Taq DNA Polymerase High Fidelity kit (Invitrogen, #11304011) and specific primers. PCR products were electrophoresed and visualized in GelRed Nucleic Acid (Biotium, Fremont, CA, USA) stained gels (1.8% agarose) under UV light. The resulting bands were purified using QIAquick ® Gel Extraction kit (Sigma-Aldrich) following manufactures instructions and sequenced by Sanger. Primers will be provided upon request.

### Statistical analysis

Statistical analysis was performed using GraphPad Prism software. *In vitro* experiments were performed at least twice with biological replicates. In *in vivo* experiments a minimum of three animals was used per group. Data were represented as mean ± standard error of the mean (SEM). Normally distributed data were compared among groups using 2-tailed Student’s test. Non-normally distributed data were analyzed using the Mann-Whitney test. Statistical significance was considered according to the following P values: \**P* < 0.05, \*\**P* < 0.01, \*\*\**P* < 0.001.

## Data availability statement

Data and material would be available upon request.

## Acknowledgments

The work was supported by grants PID2019-104265RB-I00 and PID2022-137181OB-I00 funded by MICIU/AEI/10.13039/501100011033 and by “ERDF/UE” to CB; CIBERehd; Ayudas Investigador AECC 2022, INVES223049AREC to MA; FIMA predoctoral and AECC predoctoral 2025 fellowships (PRDNA258508ROJO) to CR; AEI/FSE FPI fellowship for AO (PREP2022-000614); Ramón y Cajal Program contract to MGFB (RYC2018-024475-1); Fundación Eugenio Rodríguez Pascual; Fundación Mario Losantos; Fundación M Torres and the generous donation of Mr. Eduardo Avila. We also acknowledge the technical support of the Morphology Core facility at CIMA.

## Author contributions

C.R., A.O., M.E., M.A., R.B., M.U.L. and I.U. conducted the research, provided methodology and contributed to the acquisition of data. A.G.U., L.G., A.L., and F.P. provided resources and advice. M.G.F-B. and M.A.A. contributed to interpretation of data and reviewed the manuscript. M.A. contributed to project administration and supervision. C.B. contributed to conceptualization, fund acquisition, project administration, supervision, and writing.

## Declaration of interest statement

Authors declare no conflict of interest.

## Supplementary Figures

**Figure S1.**
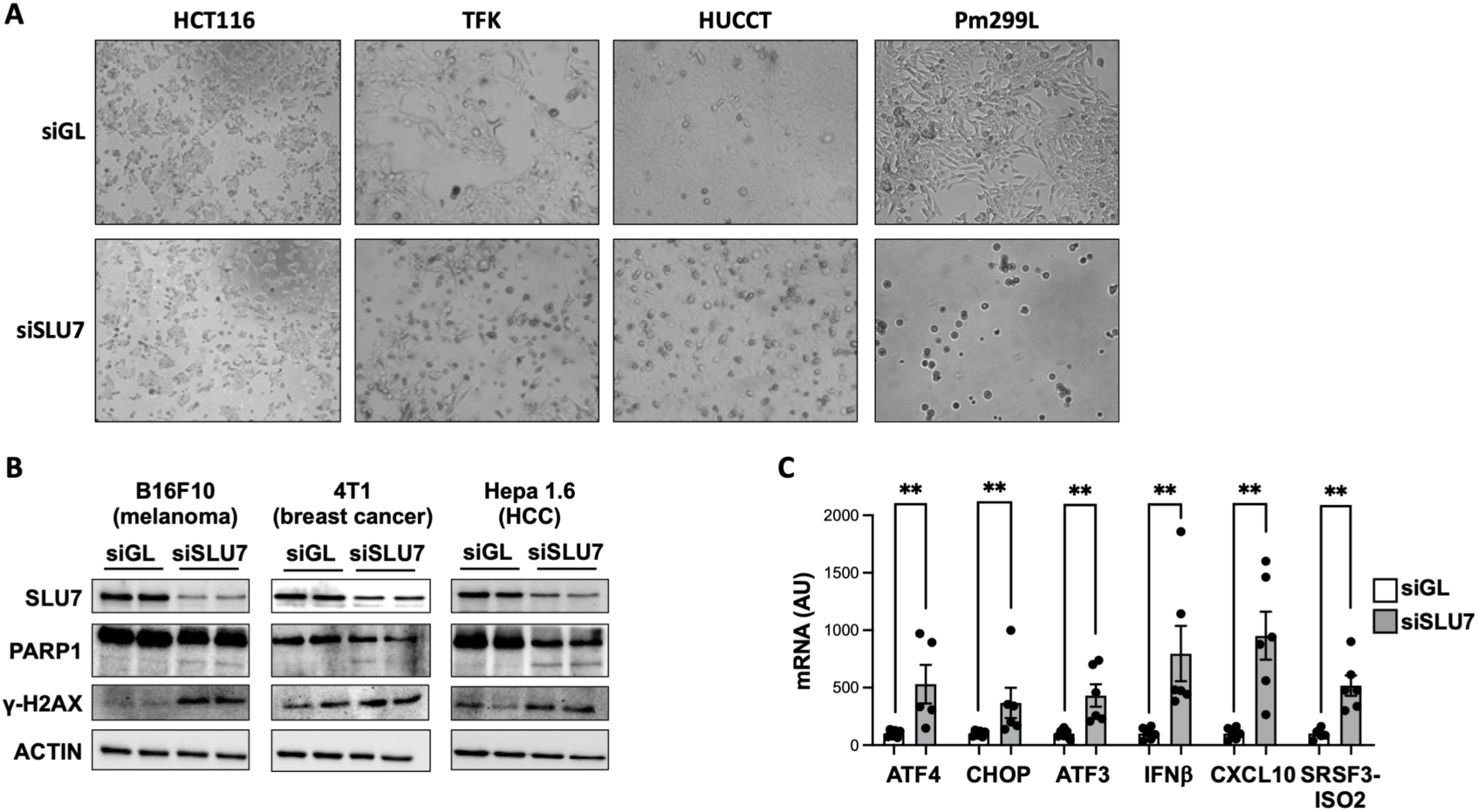
Impact of SLU7 silencing on human and mouse cancer cells viability and gene expression. **A.** Reduction of cell viability 72h after transfection with siSLU7 compared with control siGL in human HCT116, TFK-1 and HUCCT-1 cancer cells and mouse Pm299L HCC cell line. **B.** SLU7 knockdown 72 hours after siSLU7 transfection was accompanied by PARP1 cleavage and induction of γ-H2AX. The Western blot of Actin is shown as loading control. **C.** Real time PCR analysis of Pm299L demonstrated that SLU7 knockdown (siSLU7) induced the expression of stress-related genes (ATF4 and CHOP), accumulation of NMD targets (ATF3 and SRSF3-ISO2), and activation of the innate immune response (IFNβ and CXCL10).

**Figure S2.**
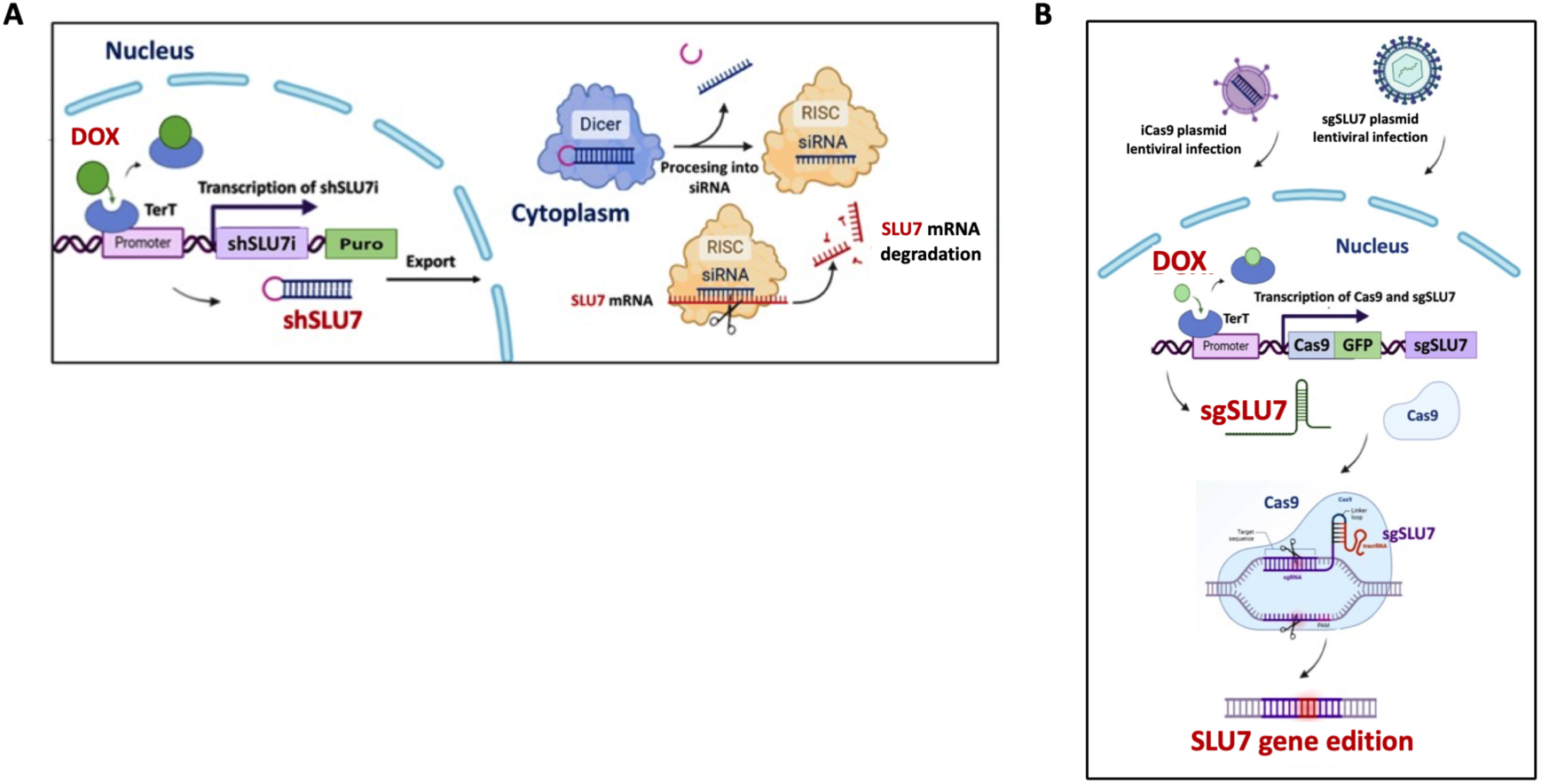
Schematic representation of the strategies used to silence SLU7 expression in an inducible manner. **A.** Scheme of the construct used to express in a doxycycline (DOX) inducible manner a specific SLU7 shRNA. **B.** Scheme of the strategy used to generate cell lines expressing in a DOX inducible manner Cas9 and constitutively sgRNAs directed against SLU7 (sgSLU7).

**Figure S3.**
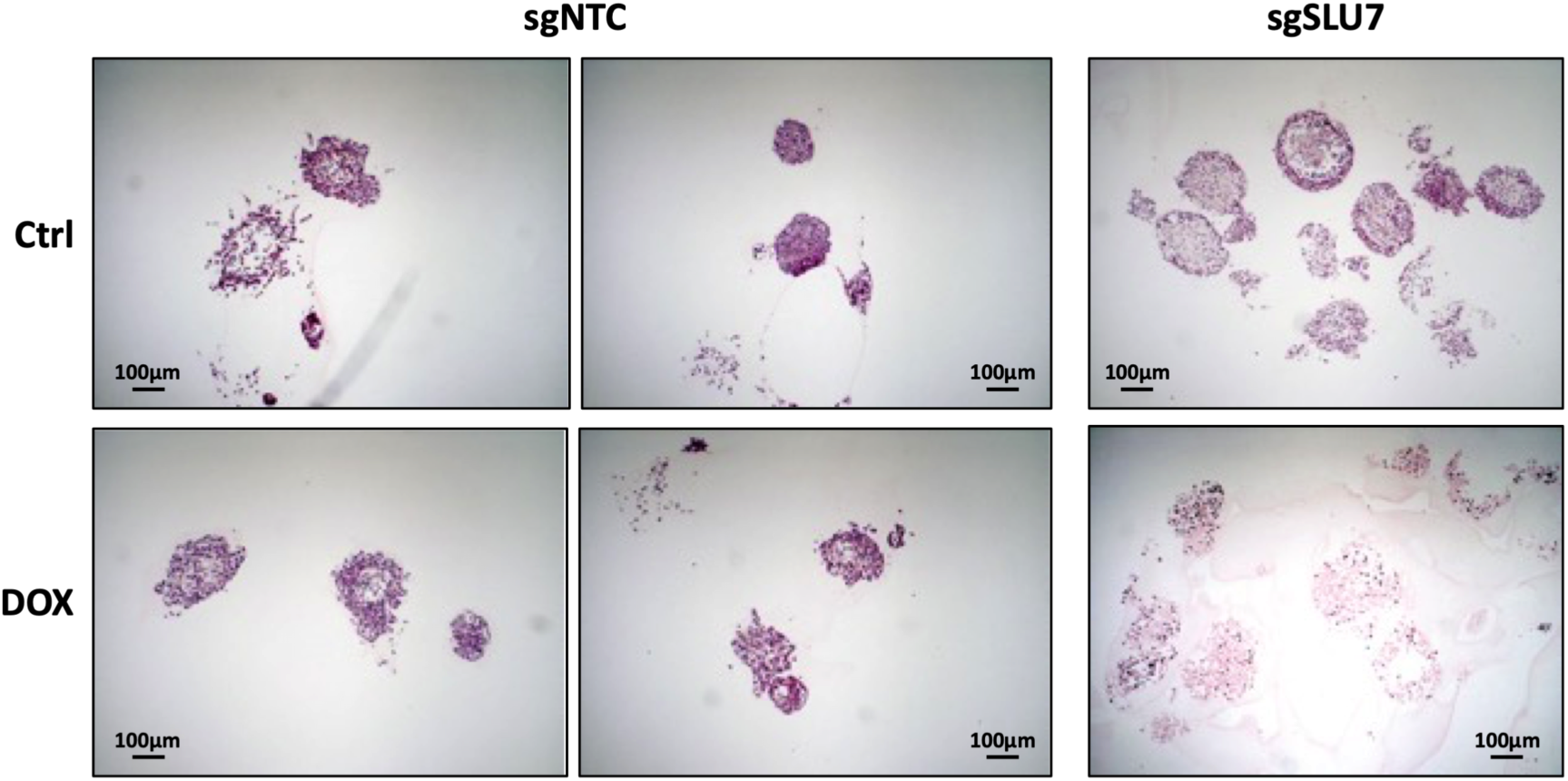
Organoids generated from liver tumors formed after intrahepatic implantation of a tissue fragment from a subcutaneous tumor previously induced with Pm299L-iCas9-sgNTC (sgNTC) or Pm299L-iCas9-sgSLU7 (sgSLU7) HCC cells. Organoids were stained with hematoxylin and eosin 72 hours after treatment with doxycycline (DOX) or vehicle (Ctrl). In sgNTC induced organoids two different fields are shown to increase the number of organoids evaluated.

**Figure S4.**
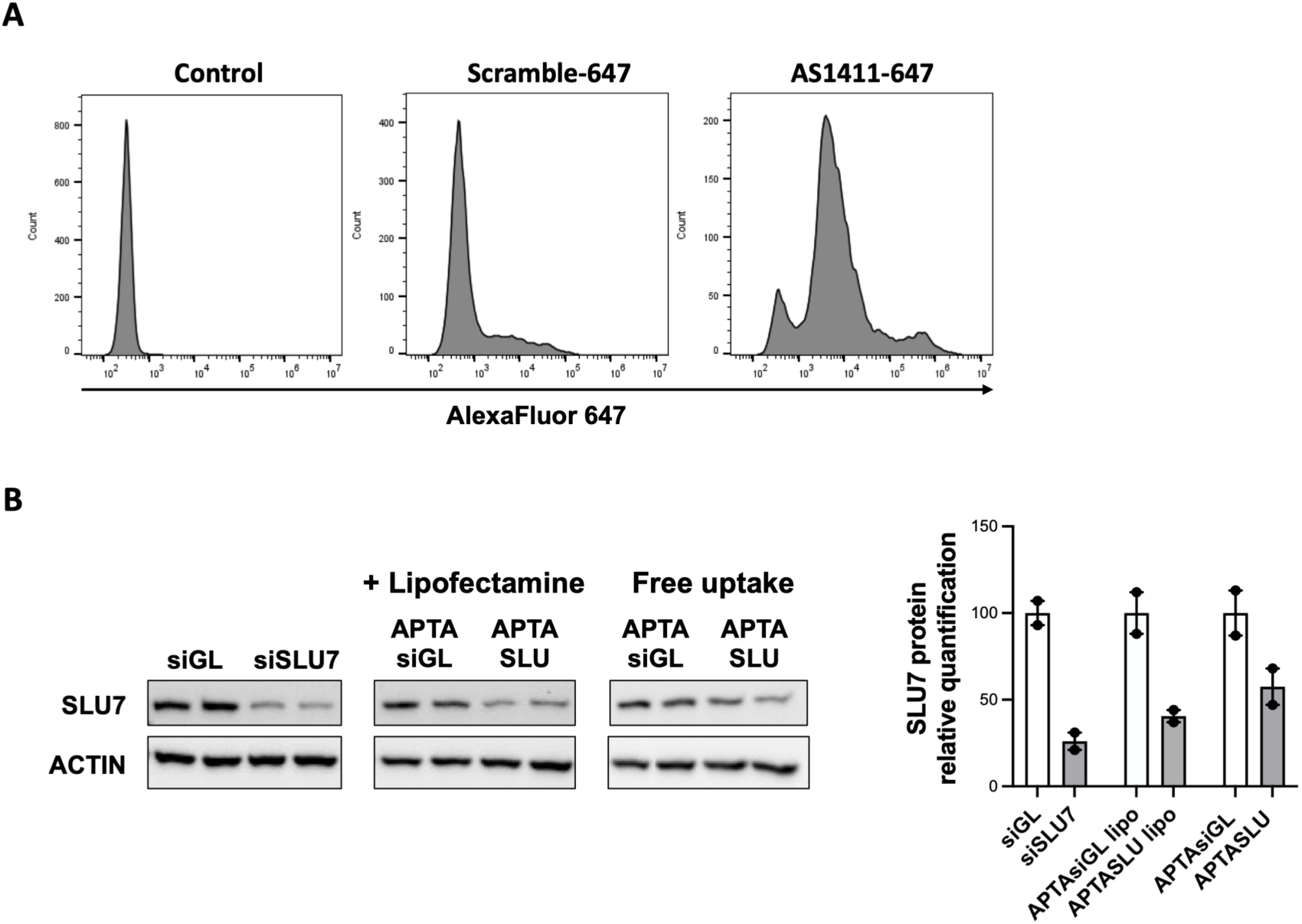
**A** Binding of AS1411 aptamer to Pm299L mouse HCC cells. The binding of AlexaFluor 647 labelled AS1411 (AS1411-647) or scramble (Scramble-647) aptamers to Pm299L cells was measured by flow cytometry. **B.** Silencing of SLU7 protein in Pm299L cells 72 hours after addition to the culture medium (free uptake) or after transfection with lipofectamine of the AS1411-siSLU7 chimeric aptamer (APTASLU) or control chimera (APTAsiGL). Histograms represent the quantification.

**Figure S5.**
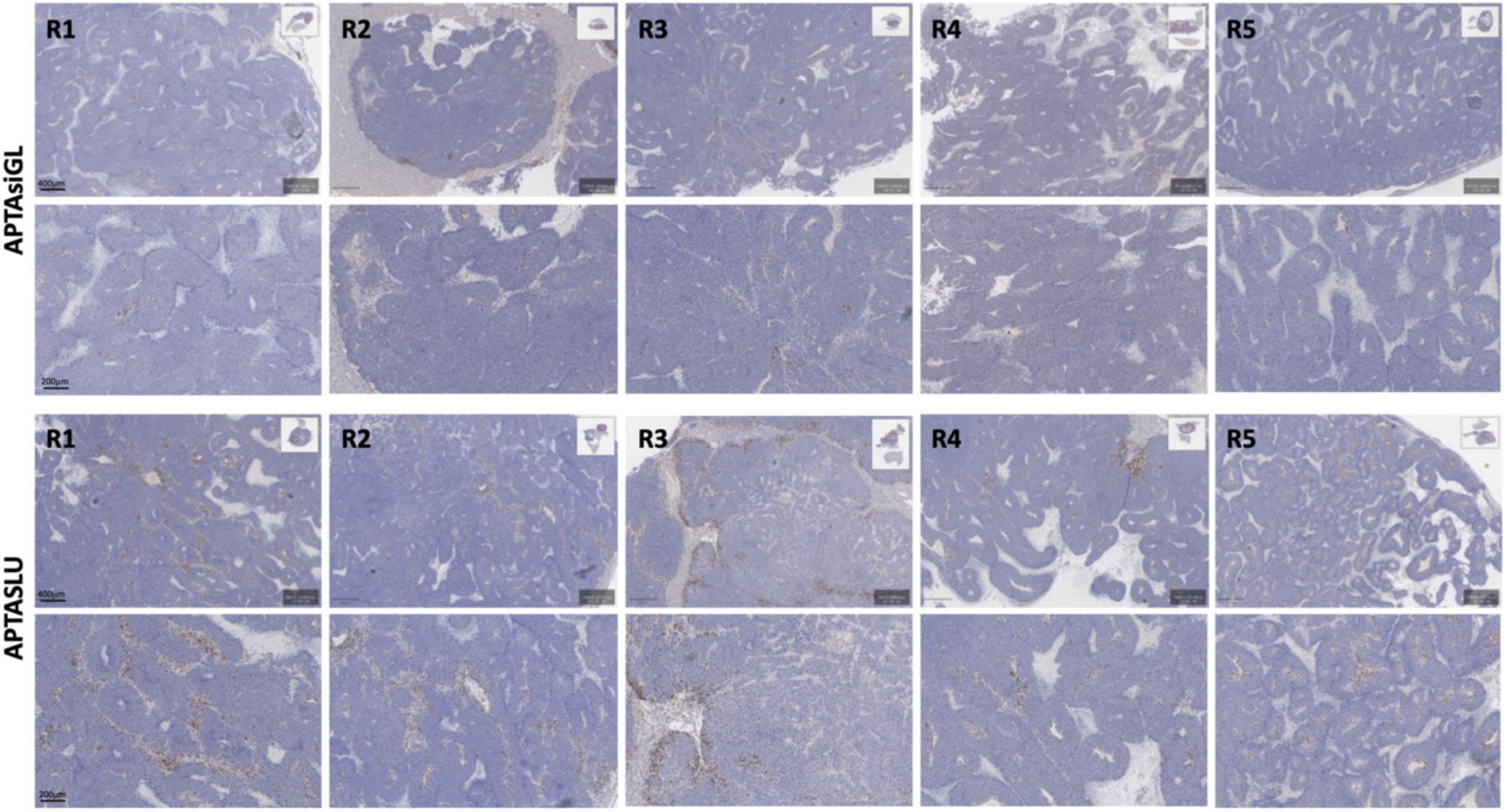
Immunohistochemical staining of CD3+ lymphocytes in liver tumors from mice treated with control (APTAsiGL) or APTASLU chimeric aptamers. Representative tumor images of 5 mice per group are shown at two different magnifications (400 µm and 200 µm). Images were obtained with QuPath program.

## Notes

### Competing Interest Statement

The authors have declared no competing interest.

